# Phylogenetic Analysis of cpn60 UT region reveal oligonucleotide regions for the detection of prominent *Enterobacteriaceae* pathogens

**DOI:** 10.64898/2026.09.14.751583

**Authors:** Harish Babu Kolla, Kunal Singh, Ann Mary Issac, Chinmayi Mhatre, Radhika Madan Urs, Joseph Kingston

## Abstract

The *Enterobacteriaceae* pathogens like *E. coli, Shigella, Klebsiella* and *Salmonella* cause a wide range of gastrointestinal and other mucosal infections when the contaminated food and water are consumed. Conventional microbiological methods are time-consuming and laborious process to detect these pathogens. Molecular assays are fast and sensitive to detect the pathogens. But a potential target for the detection is very essential to develop a molecular assay for the rapid detection. In this study, we explored the cpn60 UT region as a potential target for the detection of significant *Enterobacteriaceae* members like *E. coli, Shigella sp, Salmonella, Klebsiella* etc. We did the phylogenetic analysis of cpn60 UT region among the *Enterobacteriaceae* pathogenic groups. This analysis has revealed the presence of 13 clusters and two oligonucleotide regions with highest nucleotide conservation. Targeting these regions we have designed a set of oligonucleotide primer pairs and developed a SYBR green real time PCR assay to detect the *Enterobacteriaceae* pathogenic members. The degenerate oligonucleotide primers designed amplified a 132 bp UT region of cpn60 gene present in *Enterobacteriaceae* pathogens. The sensitivity of the assay was initially investigated by 10-fold dilution series of overnight broth culture of *E. coli* ATCC 10536 and had a detection limit of 10^1^ CFU/mL. The specificity was assessed by comparing the amplification plot of a set of *Enterobacteriaceae* pathogens and non-*Enterobacteriaceae* members like *Staphylococcus aureus* (*S. aureus*) and *Listeria monocytogenes* (*L. monocytogenes)* and found to detect all the *Enterobacteriaceae* cultures included in the study whereas amplification was not seen in *S. aureus* and *L. monocytogenes*.

## Introduction

*Enterobacteriaceae* is a large and diverse family of Gram-negative bacteria that includes several genera such as *Escherichia, Salmonella, Shigella, Klebsiella, Enterobacter, Proteus, Citrobacter*, and *Serratia* [1]. Many of these bacteria are part of the normal intestinal flora of humans and animals but can also act as opportunistic pathogens, causing gastrointestinal, systemic, and other mucosal infections through the consumption of contaminated food or water [2]. As such, members of the *Enterobacteriaceae* family are routinely monitored as indicators of hygiene and sanitary conditions [3]. Standard microbiological methods for the detection and enumeration of *Enterobacteriaceae* such as culturing on violet-red bile glucose agar (VRBGA) or enrichment in *Enterobacteriaceae* specific broths, are based on biochemical characteristics [4, 5]. However, these culture-based methods are time-consuming, often requiring 5–7 days to obtain definitive results and may lack specificity. For instance, colonies of non-*Enterobacteriaceae* such as *Aeromonas* and *Bacillus* species can also grow on VRBGA, potentially leading to false positives and misinterpretation [6]. Molecular characterization through 16S rRNA gene sequence analysis is widely used for the identification of bacteria [7, 8], but being highly conserved, precise detection of closely related genus/members is a limiting factor. On the other hand, detection by PCR followed with Sanger sequencing is laborious, expensive, time consuming and is not practical for high-throughput food testing [9, 10]. Hence, there is a need for the development of rapid, sensitive, and specific molecular assays [11–14] that can aid in Enterobacteriaceae pathogens detection. Real-time PCR (qPCR) has emerged as a preferred molecular tool for pathogen detection due to its speed, sensitivity, reproducibility, and reduced risk of cross-contamination [15, 16]. On the other hand, identification of a novel, highly specific genetic marker is also essential for improving detection. In this context, the cpn60 gene (groEL) has potential as a promising molecular marker for microbial detection [17, 18]. It encodes a 60 kDa class I chaperonin involved in protein folding and is universally present in bacteria. Though the cpn60 gene is ubiquitous in bacteria, specific detection of targeted communities can be achieved through the specific primer designing targeting the sequence variations within a region in this gene called universal target (UT) region or cpn60 UT. The cpn60 UT region is a ~500 bp long region within the gene which contains high nucleotide polymorphism to discriminate the bacterial communities. The cpn60 UT region has emerged as an alternative marker for 16 SrRNA analysis because of its sequence variability enabling the improved discrimination of closely related species [19]. This enhanced genetic resolution offers higher and precise taxonomic studies and specificity in microbial detection, especially in the case of diverse groups like *Enterobacteriaceae*. Additionally, the curated databases like cpnDB [20] offers additional support for the phylogenetic studies and complex metagenomic studies. All these features make this gene a potential molecular target for the accurate microbial detection. Moreover, the utility of this region has been demonstrated in various microbial identification studies, including differentiation of toxinotypes in *Clostridium perfringens* and detection of *Campylobacter* species [21, 22]. Till date, cpn60 has not been reported in the molecular detection of Enterobacteriaceae pathogens [19] [21–26]. The novelty of this study is employing the cpn60 UT region for the detection of significant *Enterobacteriaceae* pathogens. The present study aims to design a set of novel degenerate primers targeting conserved regions within the cpn60 UT to develop a SYBR Green-based real-time PCR assay. The primers were optimized using a SYBR green real-time PCR assay and used to detect *Enterobacteriaceae* vs non-*Enterobacteriaceae* pathogens. We have also evaluated the assay’s sensitivity using serial dilutions of bacterial cultures. This approach aims to provide a rapid and reliable tool for the detection of *Enterobacteriaceae* pathogenic members in diagnostic laboratory settings.

## Materials and methods

### Bacterial strains

The bacterial strains used in the current study (Table 1) were from the in-house bacterial collection at Defence Institute of Biodefence Technologies (DIBT), Mysore, Karnataka, India. All the bacterial strains were confirmed biochemically using BD Phoenix™ M50 (BD Diagnostic systems, MD, USA) before starting the study. The glycerol stocks of each strain were maintained upon biochemical confirmation. All bacteria were cultured in Brain Heart Infusion broth (Himedia, India) at 37°C incubator with constant shaking.

### Sequence retrieval and Multiple Sequence Alignment

For the identification of a conserved region in the cpn60 UT region, we did multiple sequence alignment among the *Enterobacteriaceae* pathogens. For this, a total of 236 nucleotide sequences of cpn60 UT region were retrieved from the genomes of the common *Enterobacteriaceae* pathogens. Corresponding nucleotide sequences of cpn60 UT region were retrieved from cpnDB (http://www.cpndb.ca/) database which includes sequences of chaperonin proteins or heat shock proteins20. Multiple sequence alignment for these sequences was performed with ClustalW module of MEGA-X software using the default parameters in the software. Gap opening penalty and Gap extension penalty for the alignment were 15 and 6.66 respectively. ClustalW was used as DNA weight matrix to perform multiple sequence alignment. Transition weight and delay divergent cut off percentage were set at 0.50 and 30% respectively. Negative matrix was turned off during the alignment.

### Phylogenetic analysis

Later, we did the phylogenetic analysis to determine the degree of evolutionary conservation and distance within the considered *Enterobacteriaceae* pathogens. Phylogenetic tree for the obtained alignment file was constructed by Neighbor-Joining (NJ) method using MEGA-X software. Phylogenetic tree was constructed with 1000 bootstrap replicates by the modified Nei-Gojobori (Jukes cantor) model based on the standard genetic code using synonymous-Nonsynonymous substitution parameter. Only synonymous substitutions were included with transition/transversion ratio 2.00. Polymorphism among the *Enterobacteriaceae* members used in the study was also analyzed with MEGA-X software and polymorphism among the sequences in the alignment file was identified. Conserved (C), Variable (V), Parsim-info sites (Pi), Singletons (S), Tajima’s neutrality statis (D), transition-transversion ratio (R), ps, Θ were calculated from the MEGA alignment file. After the phylogenetic analysis, the degenerate primers were designed based on the conservation of nucleotides within the clusters formed in the phylogenetic tree.

### Degenerate primer designing

For the design of a set of degenerate primers, we first identified a possible conserved regions within a range of an amplicon. We later targeted those regions for the primer design. The nucleotide variations at these two regions (one coding for the forward and the other for reverse primer regions) were highlighted and the nucleotide degeneracy was brought at these sites in all the phylogenetic clusters in *Enterobacteriaceae* pathogens. For examples, if a nucleotide site is ‘A’ in the cluster I and ‘G’ in the cluster II, that site is coded as ‘R’ in the primer design that codes for both the A and G enabling the detection of both the clusters. We followed a similar strategy for designing the degenerate primers as per the universal nucleotide degeneracy code of IUPAC. Oligonucleotide primers with degenerate nucleotide positions were designed for the detection of *Enterobacteriaceae* cultures mentioned in Table 1.

### Real-time PCR Assay

The SYBR-green based real-time PCR assay was optimized using *E. coli* ATCC 10536 and *S. aureus* ATCC 700699 (negative control). The reaction was carried out in a 10 μl final volume containing 5 μl of 2X TB green Premix Ex-Taq master mix (Takara), 2 μl of degenerate forward and reverse primers (5 pmol), 0.2 μl of ROX reference dye (50X), 1 μl of boil lysate template (108CFU/mL) and 1.8 μl of Milli-Q water [27]. The reactions were performed with a thermal cycling condition of 40 cycles at 95°C for 30 sec, 95°C for 5 sec, 60°C for 20 sec followed by a melt curve stage in QuantStudio 3 thermal cycler system (Applied Biosystems). ROX was used as a passive internal reference dye in the real-time PCR assay.

### Sensitivity

To determine the sensitivity of the SYBR green based real-time PCR assay, standard curves were generated from a serial 10-fold dilution of thermal lysate (108 CFU/mL to 10^1^ CFU/mL cells) of *Enterobacteriaceae* member *E. coli* ATCC 10536. The bacterial culture dilutions were used for DNA extraction by thermal lysis method. Briefly, 1 mL of bacterial culture was pelleted at 10,000 rpm for 1 minute and the supernatant was removed. The pellet was resuspended in 100uL MilliQ water and kept in a heating bath for 90°C for 15 minutes. After heating, the tube was centrifuged and the supernatant, i.e. thermal lysate, was removed to a new tube which was used as template for real-time PCR. The real-time PCR reaction was amplified in duplicates as per our previous reports and others [28].

### Specificity

The specificity of the degenerate primer pair was determined by using pathogenic *Enterobacteriaceae* members and non-*Enterobacteriaceae* bacterial strains like *S. aureus* and *L. monocytogenes* **(Table 1)**. All specificity tests were performed in duplicate using thermal lysates from 108 CFU/mL bacterial cultures as template.

### Data analysis

The real-time PCR assay was performed in duplicate. Data analyses were performed using QuantStudio Design and Analysis software v 1.4.3 of Quant Studio 3 Real-time PCR System.

## Results

### cpn60 UT as a diagnostic marker for *Enterobacteriaceae* pathogens

From the bioinformatics analysis, the cpn60 UT region of *Enterobacteriaceae* was found to have a total of 343 conserved bases and 212 variable bases **(Table 2)**. Phylogenetic analysis of cpn60 UT region revealed the presence of 13 clusters in the *Enterobacteriaceae* pathogenic members **(Supplementary figure 1)**. Similarly, the average pairwise genetic distance within and between the groups/clusters were calculated. Clusters I and V in the phylogenetic tree exhibited a lowest intra-cluster genetic distance, each with an average distance (D) of 0.01 and a standard error (SE) of 0.00, indicating a high degree of sequence similarity and suggesting that these groups represent closely related strains or lineages. Similarly, Cluster XII showed low diversity (D = 0.02, SE = 0.00), while Clusters IX (D = 0.03, SE = 0.00), VIII (D = 0.04, SE = 0.00), and XI (D= 0.04, SE = 0.01) demonstrated moderate levels of genetic variation among members (Table 3). In contrast, the clusters XIII and X showed relatively high intra-cluster genetic distances, with average values of 0.05 (SE = 0.01) and 0.06 (SE = 0.01), respectively **(Table 3)**. This suggests that these clusters comprise more genetically diverse sequences, potentially indicating the presence of multiple sub lineages or a broader evolutionary divergence within these groups. Overall, the intra-cluster distance analysis indicates varying levels of genetic relatedness across the identified clusters, ranging from highly conserved (Clusters I and V) to moderately and highly diverse (Clusters X and XIII). These findings provide insight into the evolutionary structure of the analyzed taxa and may inform further investigation into their taxonomic or functional relationships.

The average pairwise distances between the clusters ranged from 0.0083 to 0.2638, indicating variable levels of divergence among the groups. The lowest inter-cluster distance was observed between Cluster I and Cluster II (0.0083), suggesting close evolutionary relatedness. In contrast, the highest distance was recorded between Cluster XIII and Cluster VIII (0.2638), reflecting substantial divergence between these groups **(Table 4)**.

Overall, clusters formed discrete groups with intra-group distances generally lower than inter group distances. Notably, clusters I, II, IV, and III showed relatively lower mean distances among themselves, forming a tighter phylogenetic group. Conversely, clusters XII, XIII, and VIII were more distantly related to other clusters, indicating greater genetic diversity within these groups. These distance patterns support the phylogenetic clustering observed and highlight both closely and distantly related taxa within the dataset offering a high chance for the location of conserved nucleotide sequences for the primer design.

### Degenerate primer design

Two conserved regions i.e. 225-245 (Forward primer region) and 337-356 (Reverse primer region) in the aligned sequences with least number of variable nucleotides were identified and degeneracy was brought at these segregating sites according to the universal nucleotide degeneracy code. Briefly, the nucleotide positions 227, 234 and 237 in the forward primer regions contain all the four nucleotides and hence these positions are designated to be N which codes for any of the four nucleotides **(Figure 1A)**. Similarly for the reverse primer design, the nucleotide position 345 as Y coding T or C, 348 and 351 as R coding A and G and the N for 354th site **(Figure 1B)**. The selected reverse primer region is reverse translated to design a reverse primer as shown in the **Table 5**.

**Figure 1.**
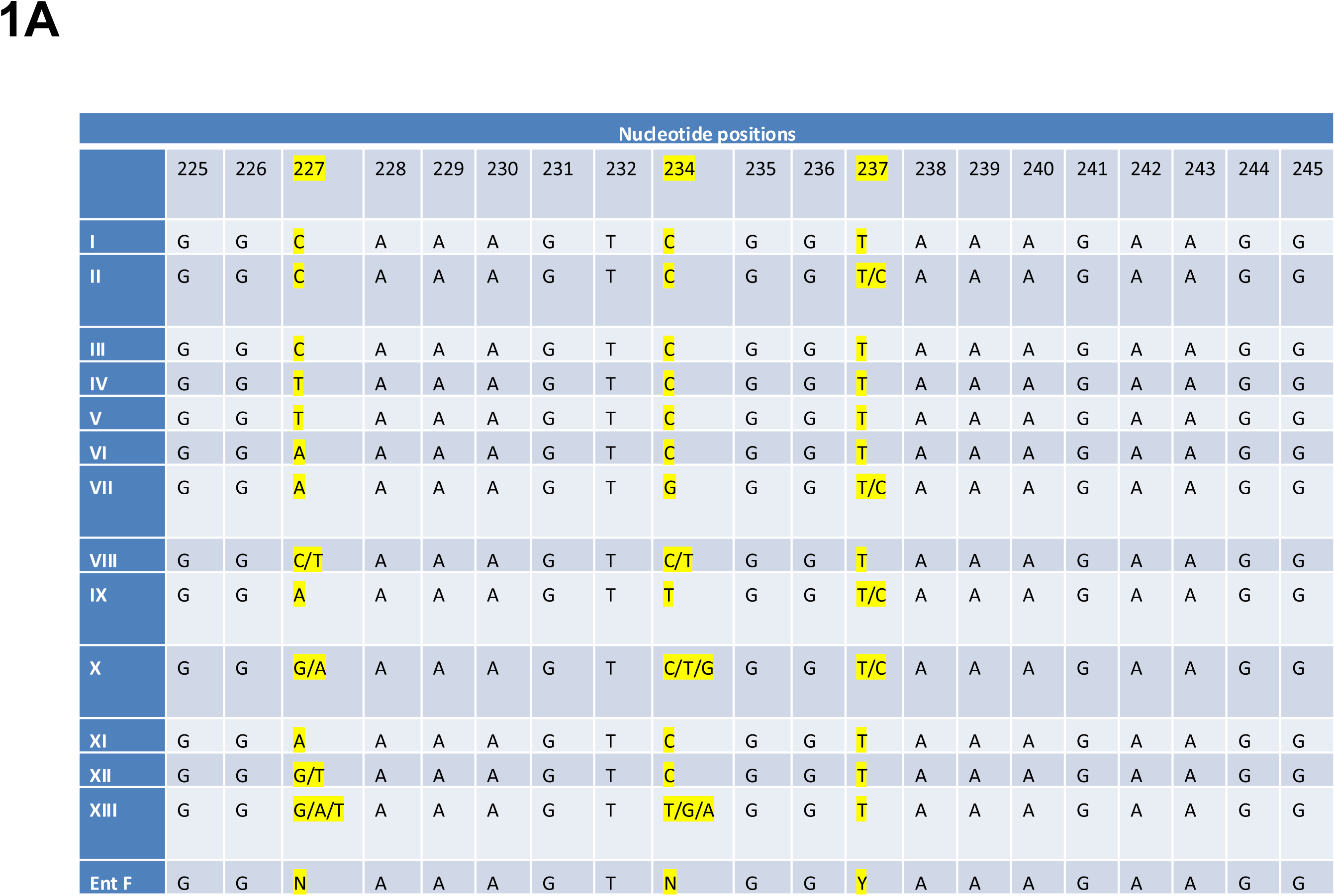
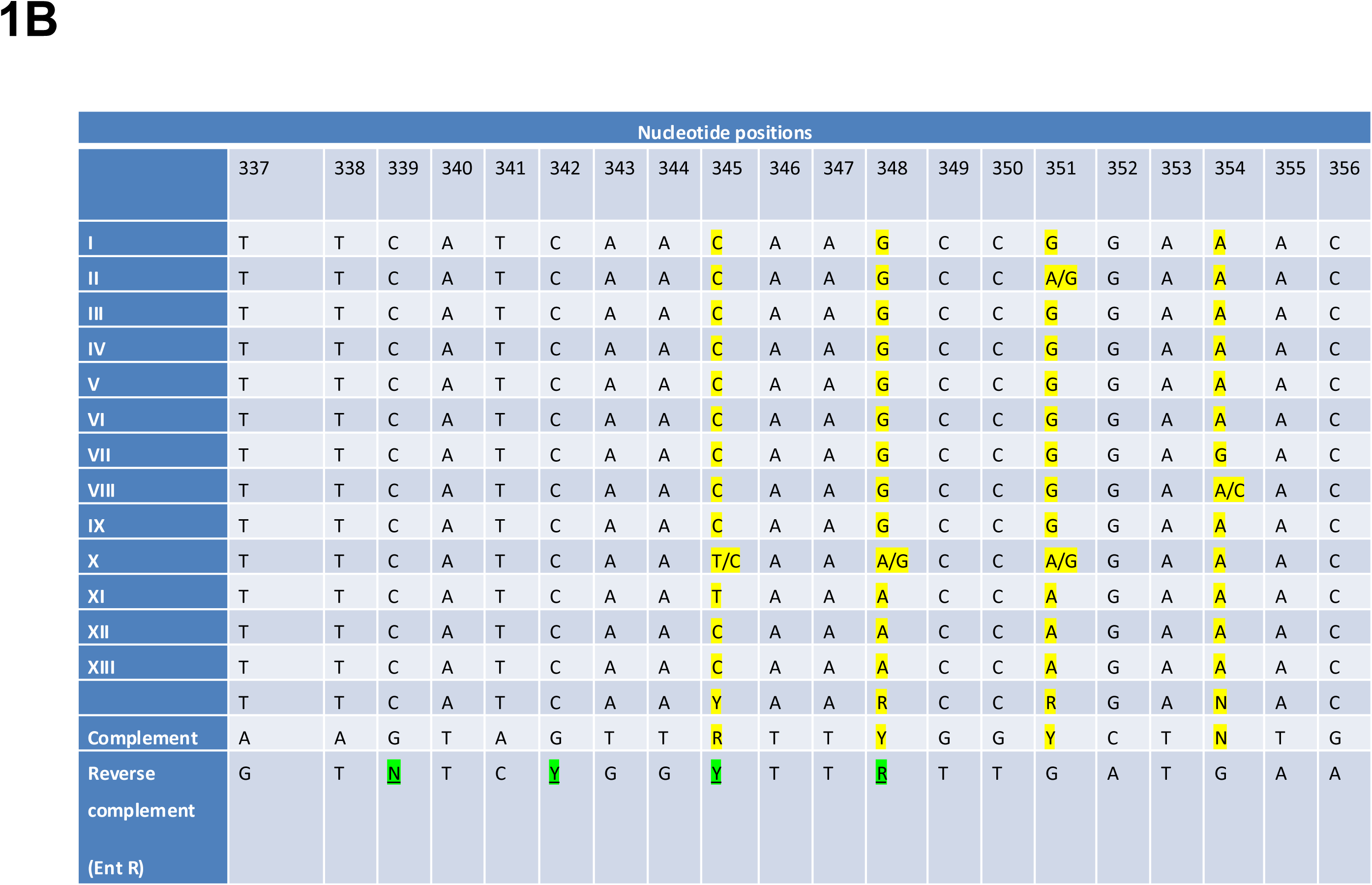
Nucleotide regions in the cpn60UT region of *Enterobacteriaceae* pathogens for **A**. The design of degenerate forward primer and **B**. reverse primers. The nucleotide positions highlighted in yellow are the positions with nucleotide variations. Degeneracy was brought at these sites following the universal nucleotide degeneracy code designated by IUPAC.

### Real-time PCR assay

The real-time PCR assay targeting cpn60 gene target was optimized with the DNA isolated from the thermal lysis from *E. coli* and *S. aureus* cultures following the real time PCR conditions mentioned in Section 2.5 and 2.6. The purity of DNA was measured using nanodrop spectrophotometer. The A260/280 ratio was in the range of 1.8-2.0. E. coli ATCC 10536 was amplified after the 13th cycle in the real-time PCR with a Ct value of 13, whereas S. aureus remained undetected **(Figure 2A)**. No template control (NTC) also showed no amplification and remained undetected. The melt curve plot indicated a single melt peak corresponding to 84.5 °C for *E. coli* ATCC 10536 indicating the presence of a single PCR amplicon in the real time PCR assay **(Figure 2A)**.

**Figure 2.**
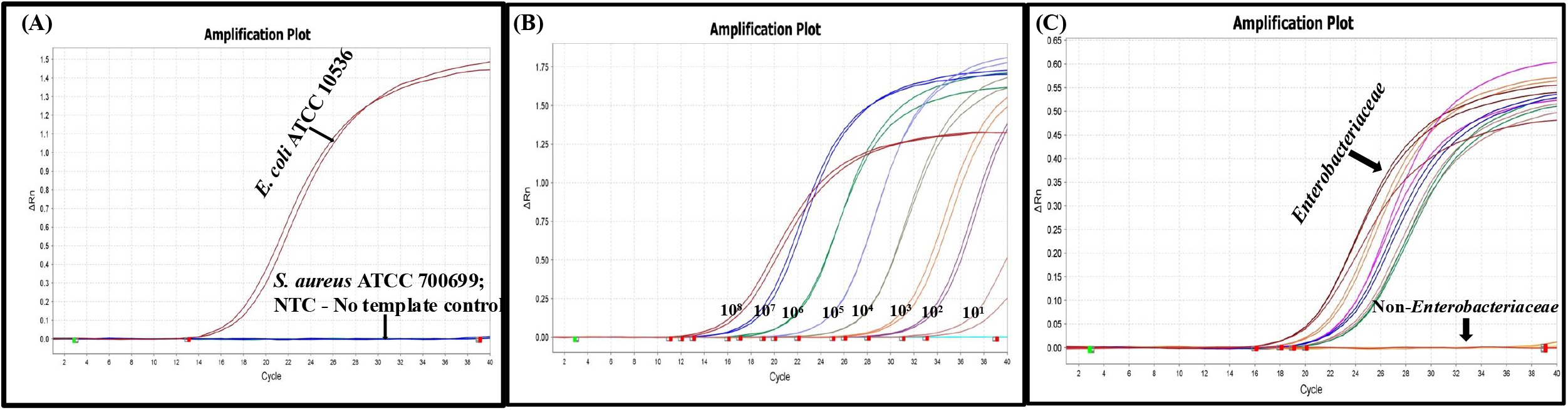
SYBR green based real-time PCR for *Enterobacteriaceae* pathogens. **A**. Amplification plot with *E. coli* ATCC 10536 (108 Cfu/mL) obtained by SYBR green based realtime PCR. **B**. The amplification plot of 10-fold serial dilution of E. coli ATCC 10536 culture (108-101 serial dilution) in SYBR green PCR assay. **C**. The amplification plot of real time PCR assay involving a significant pathogenic member of *Enterobacteriaceae* and non-*Enterobacteriaceae* like *S. aureus* and *L. monocytogenes* respectively.

### Sensitivity and Specificity

The sensitivity or limit of detection of the assay was investigated using serially 10-fold diluted *E. coli* ATCC 10536 from 108 – 10 CFU/mL using the standard protocols we followed previously and as per the previous reports27-29. The Ct values for the real-time PCR ranged from 13 to 34.5 for a culture concentration range of 10^8^-10 CFU/mL. The SYBR green real-time PCR assay had a detection limit of 101 CFU/mL (Figure 2B). The specificity of the assay was determined in both *Enterobacteriaceae* and non-*Enterobacteriaceae* cultures. All the pathogenic strains of *Enterobacteriaceae* were detected in the real-time PCR with a Ct value range of 18-22 while the non-*Enterobacteriaceae* strains (*S. aureus* and *L. monocytogenes*) remained undetected indicating the specificity of our assay to the list of cultures used in the study **(Figure 2C) (Table 1 and 6)**.

## Discussion

Detection of *Enterobacteriaceae* pathogens is significant in diagnostic microbiology as these pathogens directly contribute for the gastrointestinal and other mucosal infections. Traditional detection of *Enterobacteriaceae* pathogens involve multiple steps including plating in media such as *Enterobacteriaceae* Enrichment (EE) agar and violet-red bile glucose agar (VRBGA) which can take 5-7 days for a definitive result. Alternative rapid DNA-based methods can aid in faster detection and enumeration of *Enterobacteriaceae* family members [29, 30] where real-time PCR is used to detect and enumerate the pathogens at very low concentrations with high accuracy [28, 31–33]. Previous studies have identified protein-coding genes as suitable targets for developing diagnostic assays for the detection of bacterial communities [34]. Among them, the cpn60 universal target (UT) region within the highly conserved housekeeping gene cpn60 (also known as groEL) has been recognized for its reliability in species and subspecies-level identification, due to its higher evolutionary rate [24, 35]. This region is more polymorphic and rapidly evolving than the 16S rRNA gene and other housekeeping genes. The cpn60 gene encodes a 60 kDa class I molecular chaperonin essential for protein folding, stability, and trafficking [36]. The 549–567 bp long UT region of cpn60 could hence serve as a reliable molecular tool for bacterial identification [37]. The SYBR-green assay offers several advantages over the other molecular assays like LAMP, TaqMan-based qPCR, or 16S-ITS-based assays in terms of its easier integration in the lab workflows for the molecular analysis [38–43], lower cost since no fluorescent probes are required for the detection and easier primer design especially when employing the markers like cpn60 UT region as it provides improved taxonomic resolution for closely related species.

We specifically acknowledge the disadvantages of SYBR green assays, one being the risk of primer dimers formation and the detection of non-specific double stranded DNA [44]. To overcome these issues, we emphasized on a rigorous bioinformatics analysis and prior testing of primers in silico. Proper optimization of PCR reaction was performed using various conditions to ensure it specificity by using proper negative controls. Therefore, these measures clarify the measures taken to minimize the impact from the limitations of SYBR green based approaches. Similarly, a confidence interval for the sensitivity studies was not determined as the sensitivity of the study was performed on a limited replicates as a preliminary analytical validation rather than formal statistical modelling. However, future studies will be conducted incorporating larger replicate numbers and probit or logistic regression analysis will enable more robust estimation of LOD confidence intervals.

Furthermore, robustness of the assay was evaluated by introducing deliberate variations in key experimental parameters like DNA template concentration, primer concentration and annealing temperature. The assay demonstrated consistent amplification performance, with no cross reactivity under all tested conditions, indicating high robustness of the assay. Moreover, we also acknowledge several limitations of the current study, one such being the lack of validation of the assay on clinical and food samples. As the targeted *Enterobacteriaceae* pathogens are commonly associated with the food borne and mucosal infection outbreaks, the developed assay is evaluated directly on the DNA samples of the standard *Enterobacteriaceae* species ensuing its direct application in diagnostic laboratories. The use of cpn60 UT region as a diagnostic marker enables the discrimination from non-*Enterobacteriaceae*. Therefore, these characteristics support the suitability of the method for rapid screening applications in the diagnostics. On the other hand, the study is limited by the number of bacterial cultures used for validation. However, the primer-BLAST analysis confirmed that the designed primers exhibit high specificity form *Enterobacteriaceae* with no observed cross-reactivity with the other non-Enterobacteriaceae members (data not shown). Hence can be considered as a preliminary study to identify and validate a novel gene target to detect *Enterobacteriaceae* and further studies can be performed for additional validation in clinical and food matrices.

## Conclusion

In conclusion, the current study demonstrates the application and feasibility of cpn60 UT region as a molecular marker for the detection of *Enterobacteriaceae* pathogens through a SYBR green based real time PCR assay. High nucleotide conservation of the gene in *Enterobacteriaceae* offered a set of degenerate primers for the detection of a significant pathogenic groups in *Enterobacteriaceae* family. On the other hand, the target gene is polymorphic with the other non-*Enterobacteriaceae* members which eliminated their cross reactivity. Beyond the analytical performance, the assay offers several advantages for various practical applications such as clinal testing, food safety and beyond where the monitoring of *Enterobacteriaceae* pathogens is significant.

## Author contributions

HBK conceived the study with guidance from JK, did sequence and bioinformatics analysis, designed degenerate primers, wrote, formatted and revised the manuscript (Figure 1 and tables 2-5 and Supplementary Figure 1. KS performed the experiments (Figure 2) (Table 1 and 6), PCR data analysis and contributed to manuscript preparation. RMU, CM and AI provided the bacterial cultures, their maintenance and biochemical characterization and revised the manuscript. JK monitored the study, planned the experiments and revised the manuscript.

## Acknowledgement

All the authors thank Centre Head of Defence Institute of Biodefence Technologies, Mysore, Karnataka, India for their constant support during the study period. AI thank Defence Research and Development Organization (DRDO) for providing Senior Research Fellowship to pursue a PhD. CM thank UGC for providing Senior Research Fellowship to pursue a PhD.

## Declarations

### Conflict of Interes

The authors declare that there is no conflict of interest.

### Ethics approval

Not applicable.

### Consent to participate

Not applicable.

## Table legends

**Table 1.** List of bacterial cultures used in the study.

**Table 2.** Cpn60 UT Polymorphism among the selected Enterobacteriaceae pathogens.

**Table 3.** Average distance within the groups.

**Table 4.** Average distance between the groups.

**Table 5.** Properties of the primers designed in the study.

**Table 6.** Ct values of each strain used in the study.

