## Supplementary figures and images for "Phylogenetic Analysis of cpn60 UT region reveal oligonucleotide regions for the detection of prominent *Enterobacteriaceae* pathogens"

### Supplementary files

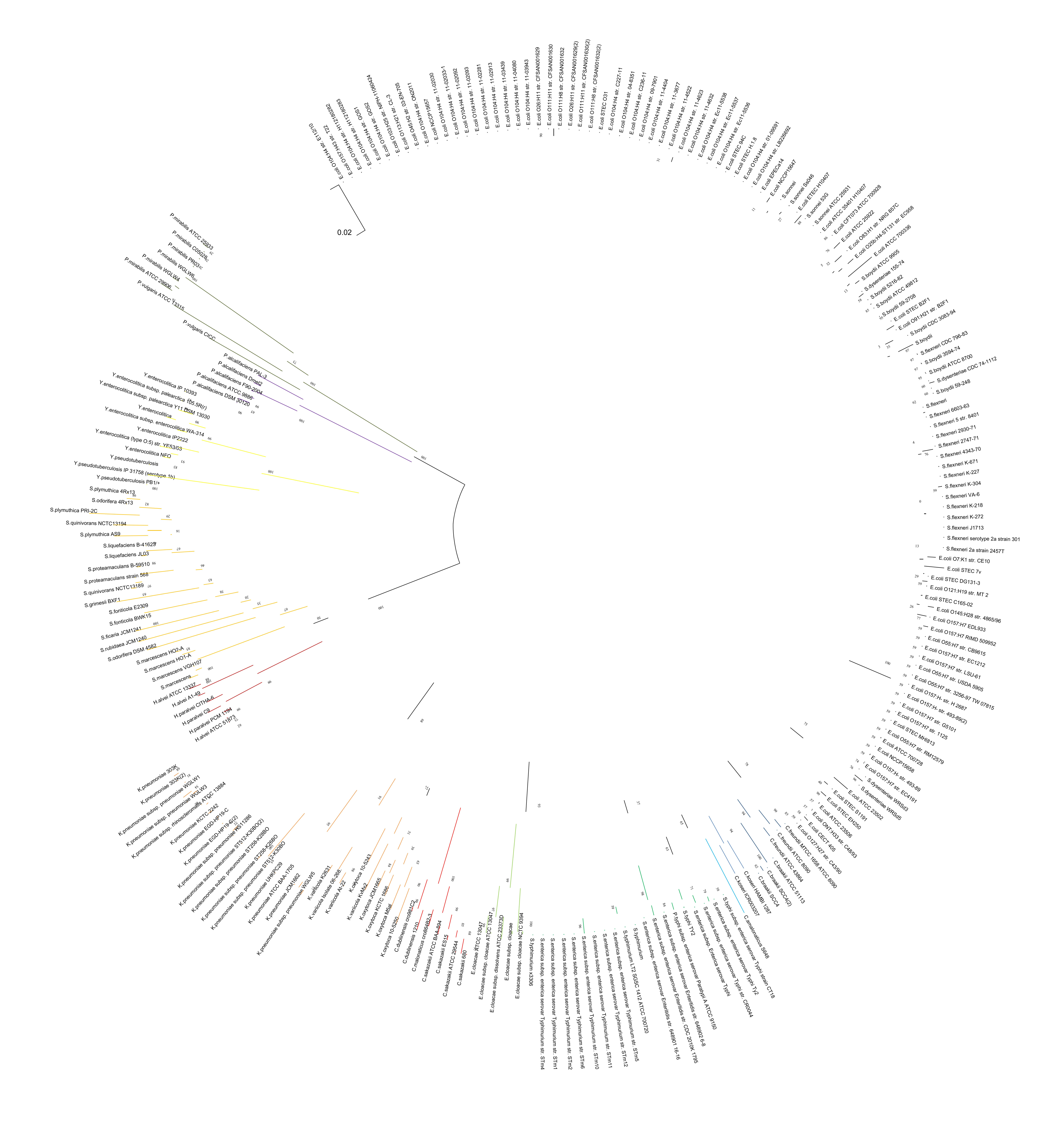
